# Isolation and characterization of autophagic bodies from yeast

**DOI:** 10.1101/2022.08.19.504482

**Authors:** Tomoko Kawamata, Shiho Makino, Yoko Kagohashi, Michiko Sasaki, Yoshinori Ohsumi

## Abstract

Autophagy is a major cellular degradation pathway that is highly conserved among eukaryotes. The identification of cargos captured by autophagosomes is critical to our understanding of the physiological significance of autophagy in cells. In the yeast *S. cerevisiae*, cells deficient in the vacuolar lipase Atg15 accumulate autophagic bodies (ABs) within the vacuole following the induction of autophagy. As ABs contain cytosolic components including proteins, RNAs, and lipids, their purification allows the identification of material targeted by autophagy for degradation. In this study, we demonstrate a method to purify intact ABs from vacuoles that retain membrane integrity and contain autophagic cargos. This technique offers a valuable tool for the identification of the cargos of autophagy, examination of autophagic cargo selectivity, and biochemical characterization of autophagosome membranes.

## Introduction

Autophagy is an evolutionally conserved intracellular degradation process in which cytoplasmic constituents and organelles are delivered to the vacuole/lysosome for degradation (1, 2). Upon induction of autophagy, a curved, membranous structure (the isolation membrane) forms, expands, and eventually forms a double-membrane structure called an autophagosome (AP) that captures a portion of the cytoplasm. The outer membrane of the AP subsequently fuses with the vacuole (or a lysosome in mammalian cells), releasing the inner membrane-bound structure into the vacuolar lumen; this structure, termed the autophagic body (AB), is then degraded by resident hydrolases. Autophagy occurs constitutively at a low level and is strongly induced when cells are starved for nutrients; in the latter case, cytosolic constituents are sequestered in an apparently random fashion into APs by macroautophagy (herein simply ‘autophagy’). Nitrogen starvation, which is sensed through the Target of rapamycin (TORC1) signaling pathway, is one of the strongest autophagy-inducing stimuli, and inhibition of TORC1 by rapamycin also induces autophagy at a level comparable to nitrogen starvation (3). Ultimately, the degradation products of autophagy are returned to the cytosol for reuse by the cell. Autophagy therefore acts as a fundamental cellular degradation pathway that maintains cellular homeostasis and forms a central plank of the starvation response. Autophagy is essential for survival during nitrogen starvation in yeast (4), and perturbations in autophagy have been associated with many diseases, including cancer and neurodegenerative pathologies in mammals (5).

AP formation is a membranous phenomenon characterized by intricate membrane dynamics. So far, 18 Atg (<u>a</u>u<u>t</u>opha<u>g</u>y-related) proteins (Atg1-10, Atg12-14, Atg16-18, Atg29, Atg31) have been identified as ‘core’ Atg proteins required for AP formation in yeast (6). Atg8 forms a conjugate with an amino group of a membrane phospholipid, phosphatidylethanolamine (PE). The formation of Atg8-PE requires Atg7 (an E1 enzyme), Atg3 (an E2 enzyme), and Atg12-Atg5-Atg16 (an E3 like enzyme). Atg4 protease functions in the processing of Atg8 precursor and delipidation of Atg8-PE. Structural and biochemical studies have recently revealed that Atg2 functions to transport phospholipids from the endoplasmic reticulum to the isolation membrane and that Atg9 acts as a phospholipid scramblase on the isolation membrane (7–11). In addition to the canonical autophagy pathway, many selective or preferential types of autophagy have been reported (12, 13). These mechanisms employ receptor and adaptor proteins to ensure the expansion of isolation membranes closely around the surface of proteins/organelles targeted by selective autophagy, indicating that the autophagy machinery can be adapted to the highly specific removal of cellular components. Whether bulk autophagy is a truly random process or exhibits some preference in cargo isolation has not yet been thoroughly investigated. A further question that remains poorly addressed in the field concerns the proteins and lipid membrane composition of AP membranes, as well as their physical properties, which so far have proven challenging to analyze due to the technical difficulties involved in specifically isolating AP membranes.

In this study, we develop a novel method for the isolation of ABs from yeast. Early electron microscopic studies of autophagy in yeast revealed that APs range in size from 300 to 900 nm in diameter and are bound by a double layer of membranes that contain very little protein, especially within the inner membrane (14). These properties of AP membranes make it challenging to purify AP by biochemical methods such as immunoisolation. In wild-type yeast cells, ABs disintegrate almost immediately within the vacuole, preventing their visualization even by live cell microscopy. In contrast, cells lacking vacuolar proteases (Pep4 and Prb1) or the vacuolar lipase (Atg15) accumulate autophagic bodies (ABs) within the vacuole (15–18). In this study, we describe a density gradient centrifugation method for the isolation of ABs from vacuoles in yeast. This approach allows for the investigation of cellular materials targeted for degradation by autophagy under diverse physiological conditions, as well as the interrogation of AB properties, such as membrane composition and biochemical features.

## Results

### Preparation of AB-containing vacuoles under autophagy-inducing conditions

To purify ABs, we started by purifying vacuoles containing ABs from cells following autophagy induction. We used *atg15*Δ cells, in which ABs accumulate within vacuoles, *atg15*Δ *atg2*Δ cells, in which APs and thus also ABs do not form, and wild-type cells, in which ABs and their cargo are degraded within vacuoles (Fig. 1A). Cells were cultured in rich media (YPD) and treated with rapamycin for 3 h to induce autophagy. Following this, spheroplasts were prepared by incubating cells with zymolyase, an enzyme that digests the yeast cell wall. Spheroplasts were then lysed by osmotic shock and vacuoles were purified from cell lysates by floatation in a Ficoll step gradient. Phase-contrast microscopy revealed that vacuoles purified from *atg15*Δ cells contain a large number of ABs, whereas vacuoles from *atg15*Δ *atg2*Δ (defective for autophagy) and wild-type (ABs degraded within the vacuole) cells are almost completely devoid of AB-like structures (Fig. 1B). Next, we examined organelle marker proteins to assess the purity of obtained vacuoles. Vacuolar membrane proteins (Pho8 and Vph1) and vacuolar luminal proteins (Pep4 and Cpy1) were enriched in purified vacuolar extracts of wild-type, *atg15*Δ, and *atg15*Δ *atg2*Δ cells compared to total cell lysates (Fig. 1C). On the other hand, mitochondrial (Por1 and Cox2), Golgi (Van1), and nuclear marker proteins (Gsp1) were largely excluded from purified vacuolar extracts, indicating the strong and specific enrichment of vacuole.

**Figure 1.**
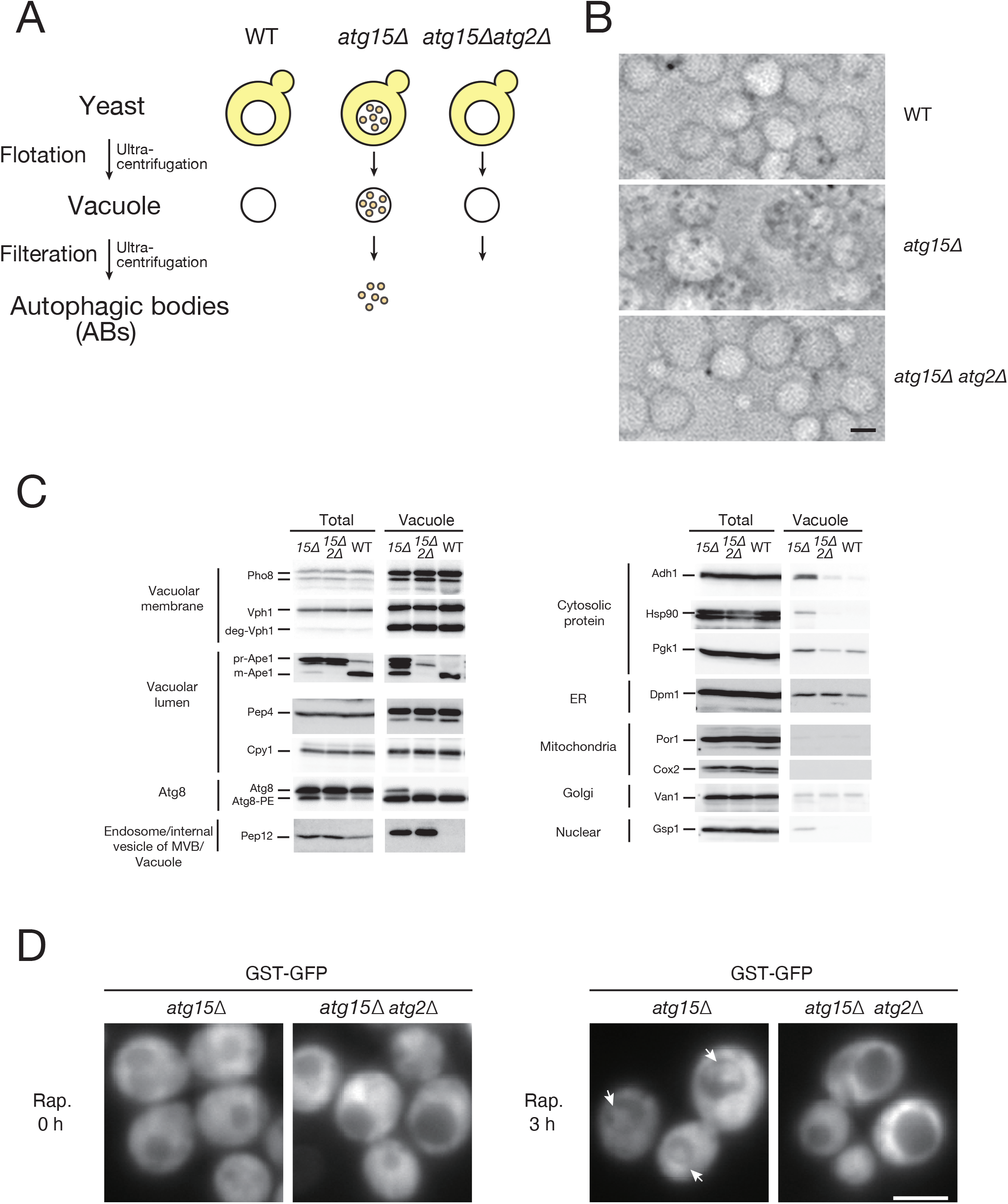
Purification of vacuole from wild-type, *atg15*Δ, and *atg15*Δ *atg2*Δ cells. **(A)** Schematic representation of the purification of autophagic bodies from vacuoles. **(B)** Phase-contrast microscopy images of vacuole fractions. After 3 hours of induction of autophagy by rapamycin, vacuoles were isolated from wild-type, *atg15*Δ, and *atg15*Δ *atg2*Δ cells. **(C)** Western blotting of the whole-cell lysate (Total) and vacuolar fraction (Vacuole) samples using antibodies against Pho8 and Vph1 (vacuole membrane), Ape1 (autophagy cargo), Pep4 and Cpy1 (vacuole lumen), Atg8 (AP and vacuole membrane), Pep12 (endosome/internal vesicle of MVB/vacuole), cytosolic protein (Adh1, Hsp90, Pgk1), Dpm1 (ER membrane), Por1 (outer mitochondrial membrane), Cox2 (mitochondrial matrix), Van1 (Golgi), and Gsp1 (nucleus). Note that Vph1 is sensitive to proteolysis and appears in part as a band with an apparent molecular mass of 75 kDa (34). (**D)** Fluorescence microscopy images of GST-GFP expressed in *atg15*Δ and *atg15*Δ *atg2*Δ cells. The localizations of GST-GFP before and 3 hours after rapamycin treatment were examined. Arrow indicates ABs.

Previous studies have suggested that the ER is a source of membranes for APs, and that most APs contain ER fragments (19). We investigated whether ER membrane proteins are detected in purified vacuolar extracts isolated from wild-type, *atg15*Δ, and *atg15*Δ *atg2*Δ cells, finding that the ER membrane protein Dpm1 is detected at a similar level in all strains (Fig. 1C). This suggests that a part of the ER may contaminate purified vacuolar fractions. We also examined Pep12, a t-SNARE that localizes mainly to endosomes/MVBs. After fusion of an MVB with the vacuole, internal vesicles of the MVB are degraded in the vacuole. It has previously been reported that *ATG15* is involved in the degradation of the internal vesicles of MVB (16). Indeed, Pep12 was detected in the vacuolar fraction of *atg15*Δ and *atg15*Δ *atg2*Δ, while no signal was detected in wild-type vacuolar fractions (Fig. 1C). The absence of a Pep12 signal in the wild-type vacuoles suggests that Pep12 was degraded in this organelle. We conclude that vacuoles purified from *atg15*Δ or *atg15*Δ *atg2*Δ cells contain internal vesicles of MVB.

### Characterization of cytosolic proteins in AB-containing vacuoles

Atg8 is a ubiquitin-like protein that is widely used as a marker of APs. It has been reported that Atg8 localizes to the vacuolar membrane independently of the autophagy machinery (20). We detected Atg8-PE in vacuole extracts from wild-type, *atg15*Δ, and *atg15*Δ *atg2*Δ cells, but a non-conjugated form of Atg8 was additionally detected in *atg15*Δ cells (Fig. 1C). The Atg8-PE observed in *atg15*Δ *atg2*Δ vacuolar fractions may arise from Atg8 bound to the outer face of the vacuolar membrane in an autophagy-independent manner. Meanwhile, free Atg8 observed only in *atg15*Δ cell extracts suggests cleavage of Atg8-PE bound to the inner side of ABs by Atg4 (See also Atg8 and Atg8-PE in Figure 2).

**Figure 2.**
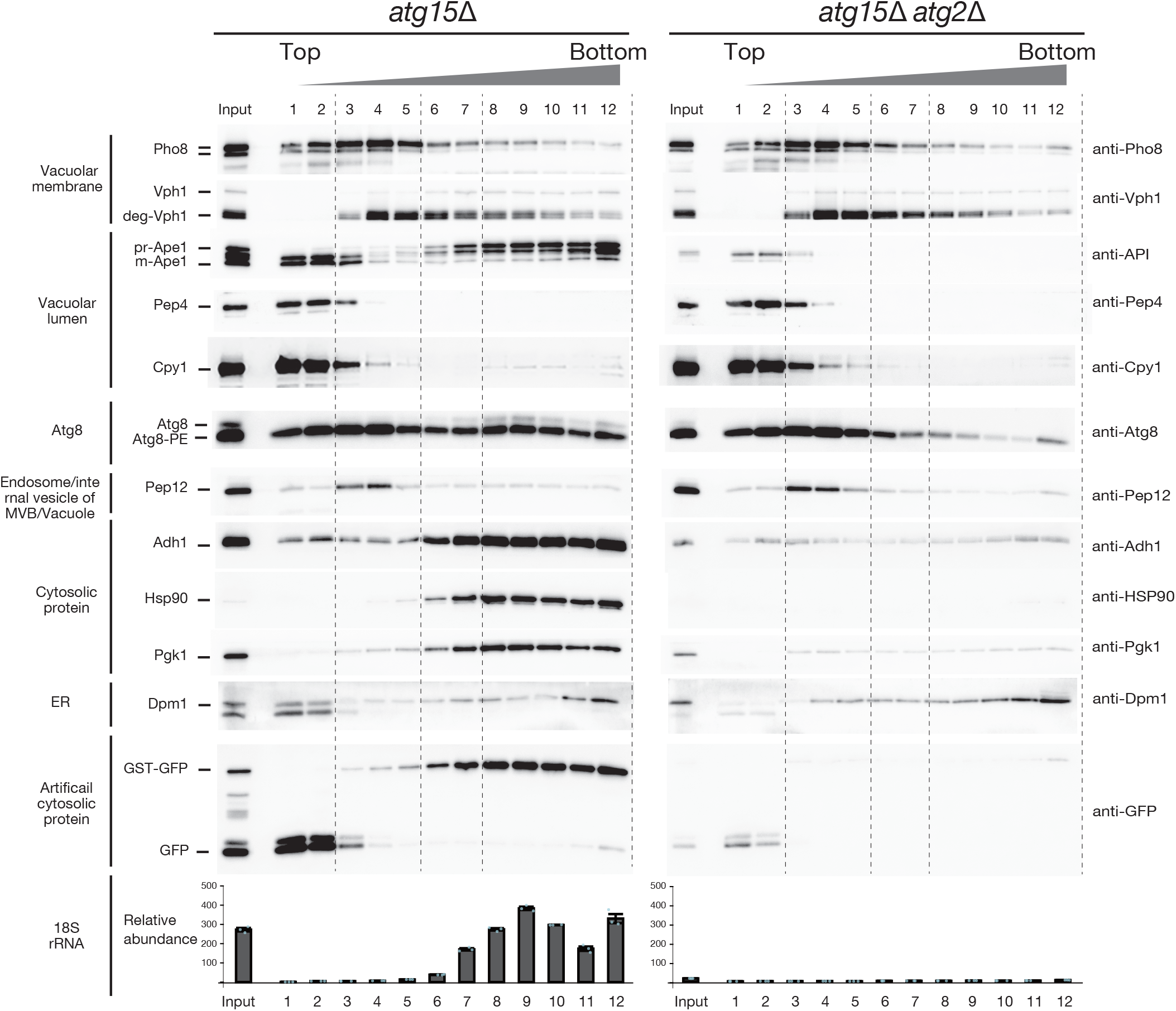
Purification of AB from *atg15*Δ cells. Separation of the AB fractions from the vacuolar lysates by OPTIPREP density gradient centrifugation. Vacuolar lysates, were passed through a membrane filter with a pore size of 0.8 μm and subjected to 0-30% OPTIPREP density gradient centrifugation. This preparation was then fractioned into 12 fractions. (Top) Western blot of marker proteins before (input) and after fractionation by OptiPrep density gradient centrifugation. Typical results are shown. (Bottom) qRT-PCR analysis of 18S rRNA. The amount of 18S rRNA was measured by qRT-PCR. RNA measurements were determined in three independent biological replicate experiments.

When autophagy is induced, the well-established autophagy cargo protein Ape1 is delivered to the vacuole by autophagy (6). prApe1 is selectively taken up by APs and transported to the vacuole where AB membranes are subsequently disintegrated. prApe1 is then processed by vacuolar proteases at the N-terminal region, yielding a mature form (mApe1). In vacuoles isolated from wild-type cells, mApe1 was detected (Fig. 1C). Meanwhile, vacuoles isolated from *atg15*Δ cells showed a prApe1 band which was not observed in *atg15*Δ *atg2*Δ vacuoles. mApe1 detected in vacuoles isolated from *atg15*Δ cells is likely due to partial degradation of prApe1 during the vacuole purification process, resulting in the cleavage of prApe1 to the mature form.

We further examined several cytosolic proteins, including Adh1 (<u>al</u>cohol <u>d</u>e<u>h</u>ydrogenase), Hsp90 (<u>h</u>eat-shock <u>p</u>rotein), and Pgk1 (3-<u>p</u>hospho<u>g</u>lycerate <u>k</u>inase). These proteins were detected in vacuolar extracts purified from *atg15*Δ cells but not wild-type or *atg15*Δ *atg2*Δ cells, indicating that these proteins reside in the AB and are degraded in wild-type cells. Taken together, these data indicate that our procedure yields vacuoles containing intact ABs (Fig. 1C).

In order to provide an additional unbiased reporter of cytosolic protein sequestration in APs, we expressed a vector-borne GST-GFP chimeric protein in cells. GST-GFP localized to the cytoplasm of cells under growing conditions. When autophagy was induced by rapamycin treatment, GST-GFP expressed in *atg15*Δ cells was delivered to the vacuole and localized to the AB, while it was not observed in *atg15*Δ *atg2*Δ cells (Fig. 1D). GST-GFP therefore provides a useful marker to assess bulk autophagy.

### Purification of ABs from *atg15*Δ cells

We next examined methods to isolate intact ABs from the isolated intact vacuoles. Neither sonication nor detergent are appropriate for this purpose. We instead reasoned specific disruption of the vacuole membrane could liberate intact ABs. To this end, we focused on the size difference between vacuoles and ABs. Vacuoles in yeast are large, usually 1-4 μm in diameter, whereas ABs are about 500 nm in diameter. Taking advantage of this difference, we attempted to break the vacuolar membrane by filtration (Fig. 1A). After passing the purified vacuoles through membrane filters of various sizes, we examined vacuoles by microscopy. We found that a 0.8 μm pore size membrane filter, which is larger than the average AB diameter, could efficiently disrupt vacuoles. To test whether ABs can be isolated intact using this approach, we isolated vacuoles from *atg15*Δ and *atg15*Δ *atg2*Δ cells expressing GST-GFP under rapamycin-treated conditions and passed these samples through 0.8 μm filters. We subjected these filtrates to OPTIPREP density gradient centrifugation and further processed these samples into 12 fractions (Fig. 2). In *atg15*Δ and *atg15*Δ *atg2*Δ cells, the vacuolar luminal proteins Pep4 and Cpy1 appeared in the top fractions (fractions 1-2). Pho8 and Vph1 (vacuolar membrane), and Pep12 (the internal vesicles of MVB) were distributed throughout the lighter fractions (fractions 3-5). Dpm1 (ER membrane) was widely distributed, gradually increasing towards the bottom fraction. Meanwhile, in *atg15*Δ cells, Ape1 showed a characteristic distribution: prApe1 was detected in heavier fractions (fractions 7-12), whereas mApe1 was identified in the top fractions (fractions 1-3). Cytosolic proteins (Adh1, Hsp90, Pgk1) fractionated into similar fractions as prApe1. However, these cytosolic markers were not detected in the heavy fraction derived from *atg15*Δ *atg2*Δ cells.

We also examined the distribution of the reporter GST-GFP construct. In *atg15*Δ samples, full-length GST-GFP was clearly observed in heavier fractions, whereas cleaved GFP was recovered in the light fractions. In *atg15*Δ *atg2*Δ samples, cleaved GFP was detected in light fractions. The distribution of Atg8 was examined along similar lines. In samples isolated from *atg15*Δ cells, the localization of Atg8-PE was widely distributed between fractions 1-12, but the localization of free Atg8 was found only in fractions 7-12. In *atg15*Δ *atg2*Δ samples, Atg8-PE was mainly found in fractions 1-5. Based on these results, we regard fractions 7-12, which showed prApe1, cytoplasmic marker protein, and free Atg8 signals, to be AB-enriched fractions.

We hypothesized that the reason why the AB fraction is denser than residual vacuolar membranes is due to the presence of ribosomes within ABs. Ribosomes are markedly dense supramolecular structures due to the large amount of RNA, and prior EM analyses indicate that most APs contain a similar density of ribosomes to the cytoplasm (14). We assessed 18S ribosomal RNA (rRNA) in each fraction by qRT-PCR. In samples extracted from *atg15*Δ cells, 18S rRNA was observed in fractions 7-12, which is consistent with the detection of cytoplasmic cargo proteins and Atg8 in these fractions (Fig. 2). 18S rRNA showed a peak at fraction 9. In contrast, *atg15*Δ *atg2*Δ samples yielded no 18S rRNA signal at all. Based on these results, we concluded that our density gradient centrifugation yields three fractions. The top two fractions are made up of soluble proteins, including vacuolar luminal proteins. Fractions 3-5 are made up of vacuolar membranes and internal vesicles of MVB. For subsequent analyses, we adopted a conservative definition of fractions 8-11 to use as the AB fraction; this is because fraction 12 is a bottom fraction that is likely to contain contaminating proteins and membranes due to non-specific aggregation. Meanwhile, fraction 7 is close to the vacuolar membrane fraction. In this way, most of the ER and inner membrane vesicles of MVB were removed from the AB fraction.

### Characterization of isolated AB from *atg15*Δ cells

We next set out to characterize ABs isolated using this method by collecting AB fractions and subjecting them to biochemical and EM analyses. First, a protease protection assay was performed to examine the membrane integrity of purified ABs. prApe1 and Hsp90 were resistant to proteinase K treatment, indicating that AB membranes remain intact (Fig. 3A). Second, AB fractions were examined by electron microscopy (EM) and compared with the vacuolar membrane fraction (Fig. 3B). In the vacuolar membrane fractions, fairly large membrane fragments were clearly observed. On the other hand, the AB membrane was hard to visualize, which reproduced our previous findings (21) and suggests that AB membranes have very low electron density. These ABs were observed to contain ribosomes, in agreement with 18S rRNA data (Fig. 2). Third, we analyzed the size of the AB fractions by dynamic light scattering (DLS). The average diameter of ABs was 379 nm, which is consistent with our EM data (Fig. 3C) and previously reported AB sizes. Finally, we compared the total protein complement purified from *atg15*Δ and *atg15*Δ *atg2*Δ cell-derived fractions. Many protein bands presumably representing diverse cytosolic proteins were detected in the AB fraction of *atg15*Δ cells (Fig. 2). In contrast, such protein bands were barely detectable in *atg15*Δ *atg2*Δ samples. We conclude that the proteins detected in the *atg15*Δ AB fraction are cargo proteins captured by APs and delivered to the vacuole.

**Figure 3.**
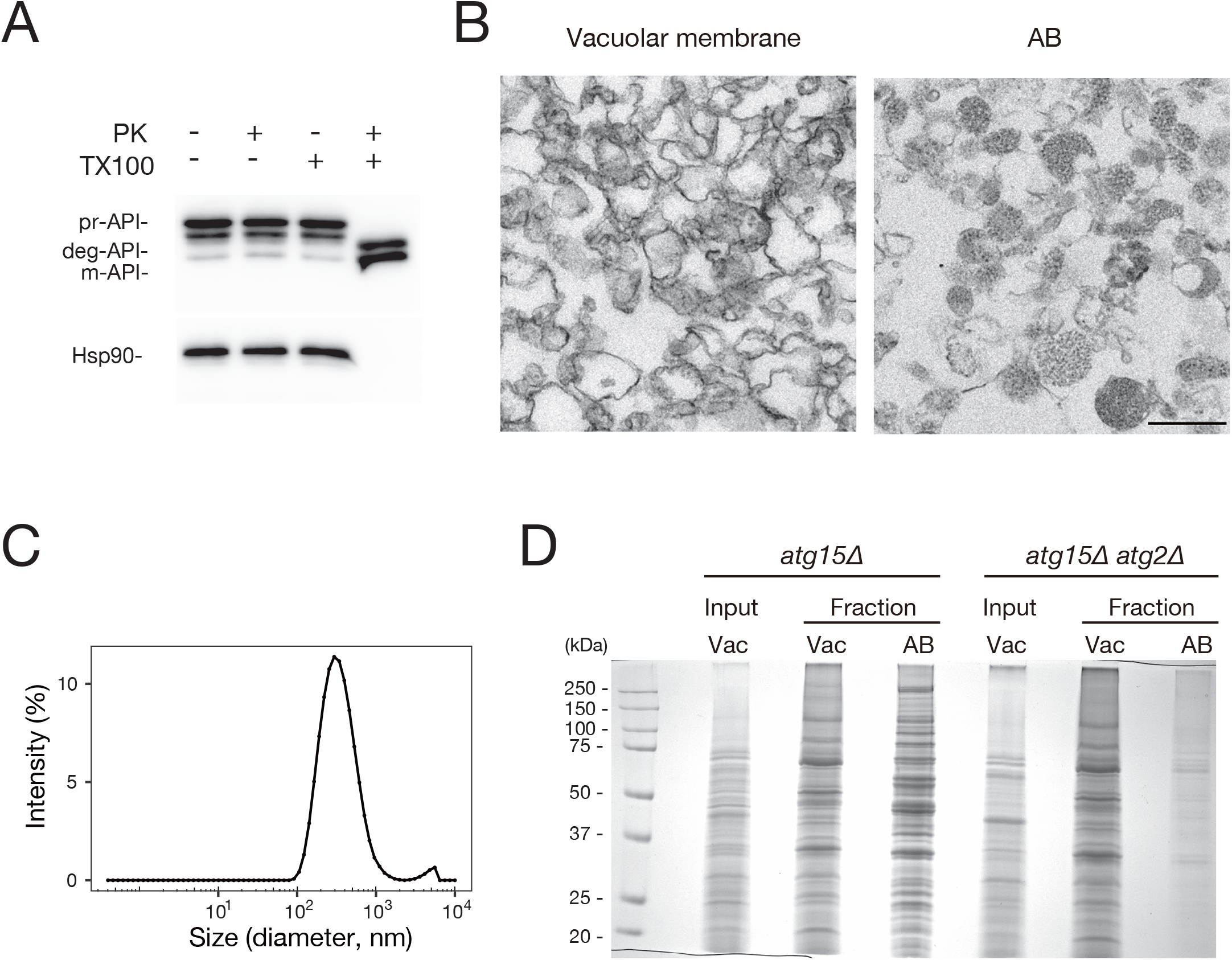
Characterization of isolated AB. **(A)** Protease protection assays after proteinase K treatment. The AB fraction was either not treated or mixed with proteinase K and/or TX-100 before being incubated on ice for 30 min. After protein precipitation, samples were analyzed by immunoblotting using anti-Ape1 or anti-Hsp90 antibodies. **(B)** Electron microscopic analyses of vacuole membrane and AB fractions. ABs typically contain cytosolic ribosomes. Scale bars, 500 nm. **(C)** Dynamic light scattering (DLS) of ABs. Before DLS, AB fractions were dialyzed to PBS. The average of four measurements for one sample is shown. The mean diameter was calculated to be 379 nm. d. nm, diameter in nanometers. **(D)** SDS-PAGE separation of proteins extracted from vacuole membrane and AB fractions of *atg15*Δ and *atg15*Δ *atg2*Δ cells. Vacuolar membrane and AB fraction exhibit distinct protein patterns when analyzed by Coomassie brilliant blue-stained SDS-PAGE.

## DISCUSSION

In this study, we present a purification method of ABs from yeast, which is the first report on the isolation of the inner membrane structure of APs. The procedure outlined in this study allows for the detailed examination of the molecular mechanisms and physiological roles of autophagy.

Our approach exploits the accumulation of ABs under autophagy-inducing conditions in the *atg15*Δ strain and the large size of budding yeast vacuoles, which are relatively easy to isolate. In order to obtain intact ABs from vacuoles, we took advantage of the size discrepancy between vacuoles and ABs, employing filtration through an empirically determined ideal pore size membrane filter to disrupt the vacuolar membrane and release ABs for subsequent fractionation. The purity of ABs was verified by western blotting of a range of representative marker proteins, and we showed that cytosolic proteins and an additionally expressed chimeric GST-GFP protein were recovered in the AB fraction in an autophagy-dependent manner. Representative mitochondrial, Golgi, and nuclear protein markers were detected only marginally, if at all, in the AB fraction. Meanwhile, Vph1, a marker of the vacuolar membrane, was observed in the AB fraction, although the majority of this protein was eliminated from ABs by density gradient centrifugation. The same was observed for ER membrane proteins, and while small amount of contamination of ER membranes cannot be ruled out, fractions 8-11 most likely include ER proteins that are naturally present in ABs due to close localization of ER or the supply of ER membranes during AP formation. Considering the overall fractionation patterns of rRNA, EM data, and SDS-PAGE analyses of total protein content, we conclude that the AB is highly enriched using the fraction strategy described in this report.

We developed this technique by purifying ABs from *atg15*Δ cells after rapamycin treatment for 3 h. We anticipate that our method is applicable to other cells that accumulate autophagic bodies, such as the *pep4*Δ *prb1*Δ strain, as well as cells treated with PMSF (15).

Purification of ABs offers a valuable and previously unavailable method for the determining and analysis of autophagy cargos. While bulk degradation via autophagy is generally described as a non-selective, “random” process, recent studies have demonstrated the selective and preferential degradation of proteins, mRNAs, and organelles by bulk autophagy (22–24). By analyzing proteins derived from purified ABs, it will be possible to directly analyze cargos captured by autophagy. This method would allow us to determine what and how much is degraded under various autophagy-inducing conditions. Furthermore, this method allows us to identify temporal changes in the complement of proteins that are degraded by autophagy.

A further implication of this study is that isolated ABs can be used to reveal characteristic features of AP membranes. A long-standing question in autophagy research concerns the origin of the AP membrane and the process of its formation. Marker proteins localized to the outer and inner membrane of APs are likely to be eliminated during the progression of AP formation and are not necessarily good markers. Purified ABs will allow for the biochemical characterization of the inner membrane of the AP, which has until now proven to be technically difficult. The identification of the membrane lipids found in AB membranes promises to provide important clues for future analysis of the origin and lipid dynamics of membranes during the formation of autophagosome. For example, the specificity of lipids carried by Atg2 and Atg9 and differences between the outer and inner membranes of APs will be elucidated. The morphological characteristic of ABs, the very thin appearance of these membranes when examined by EM, may be also accounted for by lipid analyses of AB membranes.

Characterization of purified ABs may also help researchers understand the mechanism of AB disruption within the vacuole. In wild-type cells, ABs are immediately disrupted following their delivery to the vacuole. Atg15 has been shown to act as a lipase in this process, but molecular details are only poorly understood. Using purified ABs as a substrate for in vitro assays will allow detailed characterization of AB disruption.

## EXPERIMENTAL PROCEDURES

### General methods

#### Yeast strains

Yeast strains used in this study are listed in Table 1. Strains were generated using one-step gene disruption or replacement methods as described previously (25, 26). All deletion and epitope-tagged strains constructed in this study were validated by PCR.

**Table 1.**
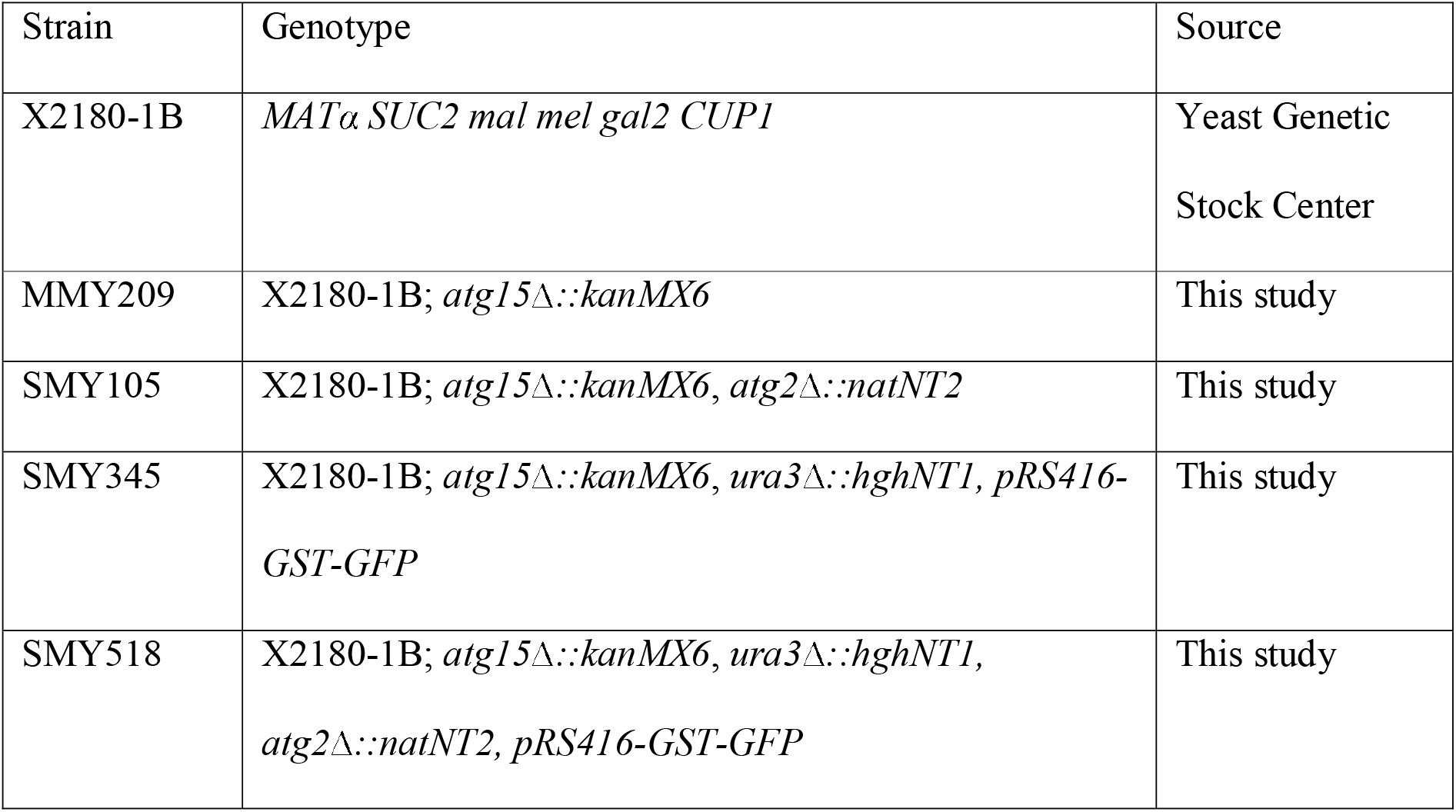
Yeast strains used in this study.

### Plasmids used in this study

#### pRS416-GPDp [URA3 CEN] GST-GFP

The GFP^*S65T*^ gene from a pFA6a-series plasmid (pFA6a-GFP-kanMX) (27) was amplified by PCR and ligated into the p416-GPD plasmid (28) between XbaI and BamHI restriction sites. Subsequently, the GST gene from the pEGKG plasmid (29) was amplified by PCR and inserted immediately upstream of GFP^*S65T*^ using the In-Fusion HD cloning kit (Takara).

### Vacuole isolation

Yeast vacuoles were isolated from whole cells as previously described (30) with some modifications. Cells grown in 2.5 liter YPD medium in a 5 liter capacity flask to a density of OD_600_ = 1.0 were treated with 0.2 μM rapamycin for 3 h. Cells were then collected and washed once with water. Spheroplasts were prepared by incubation of cells in spheroplast buffer (1.2 M sorbitol, 50 mM Tris-HCl (pH 7.5), 50 mM 2-mercaptoethanol, and 5 U/ml zymolyase 100T (Nacalai Tesque, 07665-55) for 30 min. Spheroplasts were then collected by centrifugation, washed with wash buffer (1.2 M sorbitol and 50 mM Tris-HCl pH 7.5), resuspended in 40 ml of ice-cold buffer A (0.2 M sorbitol, 12% w/v Ficoll 400, 0.1 mM MgCl_2_, 10 mM MES-Tris pH 6.9, and protease inhibitor cocktail (complete, EDTA free, Roche), and homogenized using a Dounce homogenizer on ice. All subsequent handling was performed on ice. The lysate (25 ml) was transferred to an ultracentrifuge tube and buffer B (10 ml; 0.2 M sorbitol, 8% w/v Ficoll 400, 0.1 mM MgCl_2_, and 10 mM MES-Tris pH 6.9) was layered on top. Two of these layered samples were centrifuged at 72,000 g in a P28S swinging bucket rotor (Hitachi Koki) for 30 min at 4°C. The top, white layer (a crude vacuole preparation) was collected into a new ultracentrifuge tube. This crude vacuole preparation (5 ml) was underlaid with buffer B′ (3.5 ml; 0.2 M sorbitol, 4% w/v Ficoll 400, 0.1 mM MgCl_2_, and 10 mM MES-Tris pH 6.9) and buffer C (3.5 ml; 0.2 M sorbitol, 0.1 mM MgCl_2_, and 10 mM MES-Tris pH 6.9) and subjected to a further round of centrifugation at 72,000 g in a P40ST swinging bucket rotor (Hitachi Koki) for 30 min at 4°C. The band at the 0-4% Ficoll interface containing vacuoles (approximately 500 μl ∼ 1 ml) was collected.

### OptiPrep density gradient centrifugation

Optiprep iodixanol density media was obtained from Abbott Diagnostics Technologies AS. The continuous 0-30% OptiPrep gradients were prepared in 30 mM MES-Tris (pH6.9), 0.1 M KCl, 0.5 mM MgCl_2_, 0.2 M sorbitol using a Gradient Master (BioComp Instruments, Inc.). Before loading, the vacuolar fraction was filtered (0.8 μm PC Membrane, ATTP01300, Merck Millipore) and the vacuolar fraction (less than 1 ml) was added to the top of 11 ml gradients and centrifuged at 72,000g for 90 min using a P40ST swinging bucket rotor (Hitachi Koki). Fractions were collected from the top using a Piston Gradient Fractionator (Model 152 BioComp Instruments, Inc). Twelve fractions (1 ml each for fraction 1-10, 0.6 ml for fraction 11, and 0.2 ml (bottom) for fraction 12) were collected. Aliquots (15 μl) of each fraction were resolved by SDS-PAGE and subjected to immunoblotting. For qRT-PCR analyses, aliquots (10 μl) of each fraction were used. For staining by Coomassie brilliant blue (CBB), the vacuolar fraction (640 μl) and AB fraction (640 μl) were precipitated using trichloroacetic acid (TCA) and resolved by SDS-PAGE before staining with CBB (Nacalai Tesque, 04543-64).

### Protease protection assay

AB fractions were treated with 160 μg/ml proteinase K for 30 min on ice in the presence or absence of 0.2 % TX100. Following protease treatment, the reaction was terminated by adding PMSF on ice. Samples were then precipitated by TCA and subjected to immunoblotting.

### DLS analysis

Before DLS analysis, AB fractions were pooled and dialyzed three times against PBS buffer using a dialysis membrane with a 100 kDa MWCO (Spectra/Por™ Biotech Cellulose Ester (CE) Dialysis Tube, SPECTRUM, 131414). AB in PBS was analyzed using a Zetasizer Nano S (Malvern Instruments) in 40 ul quartz cuvettes. Four measurements were performed after equilibration for 2 min at 20 □C. Experimental data were processed using the manufacturer’s software. Corrections for solvent refractive index (1.335) and viscosity (1.020) were used (Dispersion Technology Software, Malvern Instruments).

### Immunoblotting

Immunoblot analyses were performed as described previously (31–33). Primary antibodies against GFP (Roche, 11814460001), ALP (Abcam, ab113688), PGK (ThermoFisher Scientific, 459250), Vph1 (Invitrogen, A6426) and Ape1 (19), Pep12 (Invitrogen, A-21273), Dpm1 (Invitrogen, A-6429), Por1 (Invitrogen, A-6449), Cox2 (ThermoFisher Scientific, 459150), Van1 (a gift from Koji Yoda, the University of Tokyo), Gsp1 (24), Pep4, Cpy1, Hsp90, Atg8, and Adh1 (Ohsumi lab stock) were acquired from the indicated sources. Images were acquired using FUSION-FX7 (Vilber-Lourmat) imaging systems. The images were processed using the manufacturer’s software.

### Light and Fluorescence Microscopy

For isolated vacuoles, images were acquired on an inverted microscope (Olympus IX71) equipped with a 100x oil-immersion objective and phase contrast optics (UPlanFL N 100x 1.3). Intracellular localization of GST-GFP was examined using an inverted fluorescence microscope as described previously (33). Images were captured using an image acquisition system and analysis software (HCImage, Hamamatsu Photonics).

### EM analysis

The samples were fixed with an equal amount of 4% paraformaldehyde and 4% glutaraldehyde in 0.1 M phosphate buffer (pH 7.4) for 1 hour. Then they were then subjected to a second round of fixation with 2% glutaraldehyde in 0.1 M phosphate buffer (pH 7.4). After fixation, the samples were postfixed with 2% osmium tetroxide in 0.1 M phosphate. Ultrastructural analysis of vacuole membrane fraction and AB fraction was performed by Tokai-EMA Inc (Japan).

### Isolation of RNA and qRT-PCR

Isolation of RNA and qRT-PCR analysis were performed as described previously (32) (24). Briefly, RNAs from isolated vacuoles and 12 fractions yielded by OptiPrep density gradient centrifugation were extracted by TRIzol reagent (Thermo Fisher Scientific), and cDNAs were synthesized using the PrimeScript RT reagent kit with gDNA eraser (TAKARA). A random hexamer was used for cDNA synthesis. Subsequent qPCR was performed using TB Green Premix Ex Taq II (Tli RNase H Plus) (TAKARA) with the *RDN18*-specific primers 5′-AACTCACCAGGTCCAGACACAATAAGG-3′ and 5′-AAGGTCTCGTTCGTTATCGCAATTAAGC-3′. Serial dilutions of cDNA were used for qPCR calibrations. Melting-curve analyses confirmed the amplification of a single product for *RDN18*.

## ACKNOWLEDGEMENTS

We are grateful to members of the Ohsumi laboratory for discussions and critical comments on the manuscript and the Biomaterials Analysis Division, Open Facility Center, Tokyo Institute of Technology for DNA sequencing. We also thank Dr. Alexander I May for editing. This work was supported in part by Grants-in-Aid for Scientific Research 16H06375 (to Y.O.) and 18H02399 (to T.K.) from the Ministry of Education, Culture, Sports, Science and Technology of Japan.

## Author contributions

T.K. and Y.O. designed experiments and wrote the manuscript; T.K., and S.M., Y.K., and M.S. performed the experiments

The authors declare no conflicts of interest.

## References

1. Klionsky, D. J. (2007) Autophagy: from phenomenology to molecular understanding in less than a decade. Nat Rev Mol Cell Biol 8, 931–937

2. Ohsumi, Y. (2014) Historical landmarks of autophagy research. Cell Res 24, 9–23

3. Kamada, Y., Yoshino, K., Kondo, C., Kawamata, T., Oshiro, N., Yonezawa, K. and Ohsumi, Y. (2010) Tor directly controls the Atg1 kinase complex to regulate autophagy. Mol Cell Biol 30, 1049–1058

4. Tsukada, M. and Ohsumi, Y. (1993) Isolation and characterization of autophagy-defective mutants of Saccharomyces cerevisiae. FEBS Lett 333, 169–174

5. Mizushima, N., White, E. and Rubinsztein, D. C. (2021) Breakthroughs and bottlenecks in autophagy research. Trends Mol Med 27, 835–838

6. Nakatogawa, H. (2020) Mechanisms governing autophagosome biogenesis. Nat Rev Mol Cell Biol 21, 439–458

7. Maeda, S., Yamamoto, H., Kinch, L. N., Garza, C. M., Takahashi, S., Otomo, C., Grishin, N. V., Forli, S., Mizushima, N. and Otomo, T. (2020) Structure, lipid scrambling activity and role in autophagosome formation of ATG9A. Nat Struct Mol Biol 27, 1194–1201

8. Matoba, K., Kotani, T., Tsutsumi, A., Tsuji, T., Mori, T., Noshiro, D., Sugita, Y., Nomura, N., Iwata, S., Ohsumi, Y., Fujimoto, T., Nakatogawa, H., Kikkawa, M. and Noda, N. N. (2020) Atg9 is a lipid scramblase that mediates autophagosomal membrane expansion. Nat Struct Mol Biol 27, 1185–1193

9. Valverde, D. P., Yu, S., Boggavarapu, V., Kumar, N., Lees, J. A., Walz, T., Reinisch, K. M. and Melia, T. J. (2019) ATG2 transports lipids to promote autophagosome biogenesis. J Cell Biol 218, 1787–1798

10. Maeda, S., Otomo, C. and Otomo, T. (2019) The autophagic membrane tether ATG2A transfers lipids between membranes. Elife 8,

11. Osawa, T., Kotani, T., Kawaoka, T., Hirata, E., Suzuki, K., Nakatogawa, H., Ohsumi, Y. and Noda, N. N. (2019) Atg2 mediates direct lipid transfer between membranes for autophagosome formation. Nat Struct Mol Biol 26, 281–288

12. Faruk, M. O., Ichimura, Y. and Komatsu, M. (2021) Selective autophagy. Cancer Sci 112, 3972–3978

13. Suzuki, K. (2013) Selective autophagy in budding yeast. Cell Death Differ 20, 43–48

14. Baba, M., Takeshige, K., Baba, N. and Ohsumi, Y. (1994) Ultrastructural analysis of the autophagic process in yeast: detection of autophagosomes and their characterization. J Cell Biol 124, 903–913

15. Takeshige, K., Baba, M., Tsuboi, S., Noda, T. and Ohsumi, Y. (1992) Autophagy in yeast demonstrated with proteinase-deficient mutants and conditions for its induction. J Cell Biol 119, 301–311

16. Epple, U. D., Eskelinen, E. L. and Thumm, M. (2003) Intravacuolar membrane lysis in Saccharomyces cerevisiae. Does vacuolar targeting of Cvt17/Aut5p affect its function. J Biol Chem 278, 7810–7821

17. Teter, S. A., Eggerton, K. P., Scott, S. V., Kim, J., Fischer, A. M. and Klionsky, D. J. (2001) Degradation of lipid vesicles in the yeast vacuole requires function of Cvt17, a putative lipase. J Biol Chem 276, 2083–2087

18. Epple, U. D., Suriapranata, I., Eskelinen, E. L. and Thumm, M. (2001) Aut5/Cvt17p, a putative lipase essential for disintegration of autophagic bodies inside the vacuole. J Bacteriol 183, 5942–5955

19. Hamasaki, M., Noda, T. and Ohsumi, Y. (2003) The early secretory pathway contributes to autophagy in yeast. Cell Struct Funct 28, 49–54

20. Liu, X. M., Yamasaki, A., Du, X. M., Coffman, V. C., Ohsumi, Y., Nakatogawa, H., Wu, J. Q., Noda, N. N. and Du, L. L. (2018) Lipidation-independent vacuolar functions of Atg8 rely on its noncanonical interaction with a vacuole membrane protein. Elife 7,

21. Baba, M., Osumi, M. and Ohsumi, Y. (1995) Analysis of the membrane structures involved in autophagy in yeast by freeze-replica method. Cell Struct Funct 20, 465–471

22. Onodera, J. and Ohsumi, Y. (2004) Ald6p is a preferred target for autophagy in yeast, Saccharomyces cerevisiae. J Biol Chem 279, 16071–16076

23. Shpilka, T., Welter, E., Borovsky, N., Amar, N., Shimron, F., Peleg, Y. and Elazar, Z. (2015) Fatty acid synthase is preferentially degraded by autophagy upon nitrogen starvation in yeast. Proc Natl Acad Sci U S A 112, 1434–1439

24. Makino, S., Kawamata, T., Iwasaki, S. and Ohsumi, Y. (2021) Selectivity of mRNA degradation by autophagy in yeast. Nat Commun 12, 2316

25. Janke, C., Magiera, M. M., Rathfelder, N., Taxis, C., Reber, S., Maekawa, H., Moreno-Borchart, A., Doenges, G., Schwob, E., Schiebel, E. and Knop, M. (2004) A versatile toolbox for PCR-based tagging of yeast genes: new fluorescent proteins, more markers and promoter substitution cassettes. Yeast 21, 947–962

26. Knop, M., Siegers, K., Pereira, G., Zachariae, W., Winsor, B., Nasmyth, K. and Schiebel, E. (1999) Epitope tagging of yeast genes using a PCR-based strategy: more tags and improved practical routines. Yeast 15, 963–972

27. Longtine, M. S., McKenzie, A., Demarini, D. J., Shah, N. G., Wach, A., Brachat, A., Philippsen, P. and Pringle, J. R. (1998) Additional modules for versatile and economical PCR-based gene deletion and modification in Saccharomyces cerevisiae. Yeast 14, 953–961

28. Mumberg, D., Müller, R. and Funk, M. (1995) Yeast vectors for the controlled expression of heterologous proteins in different genetic backgrounds. Gene 156, 119–122

29. Mitchell, D. A., Marshall, T. K. and Deschenes, R. J. (1993) Vectors for the inducible overexpression of glutathione S-transferase fusion proteins in yeast. Yeast 9, 715–722

30. Ohsumi, Y. and Anraku, Y. (1981) Active transport of basic amino acids driven by a proton motive force in vacuolar membrane vesicles of Saccharomyces cerevisiae. J Biol Chem 256, 2079–2082

31. Kushnirov, V. V. (2000) Rapid and reliable protein extraction from yeast. Yeast 16, 857–860

32. Huang, H., Kawamata, T., Horie, T., Tsugawa, H., Nakayama, Y., Ohsumi, Y. and Fukusaki, E. (2015) Bulk RNA degradation by nitrogen starvation-induced autophagy in yeast. EMBO J 34, 154–168

33. Kawamata, T., Horie, T., Matsunami, M., Sasaki, M. and Ohsumi, Y. (2017) Zinc starvation induces autophagy in yeast. J Biol Chem 292, 8520–8530

34. Kane, P. M., Kuehn, M. C., Howald-Stevenson, I. and Stevens, T. H. (1992) Assembly and targeting of peripheral and integral membrane subunits of the yeast vacuolar H(+)-ATPase. J Biol Chem 267, 447–454

